# Homeostatic recovery of embryonic spinal activity initiated by compensatory changes in resting membrane potential

**DOI:** 10.1101/761767

**Authors:** Carlos Gonzalez-Islas, Miguel Angel Garcia-Bereguiain, Peter Wenner

## Abstract

When baseline activity in a neuronal network is modified by external challenges, a set of mechanisms is prompted to homeostatically restore activity levels. These homeostatic mechanisms are thought to be profoundly important in the maturation of the network. We have previously shown that 2-day blockade of either excitatory GABAergic or glutamatergic transmission in the living embryo transiently blocks the movements generated by spontaneous network activity (SNA) in the spinal cord. However, by 2 hours of persistent receptor blockade embryonic movements begin to recover, and by 12 hours we observe a complete homeostatic recovery *in vivo*. Compensatory changes in voltage-gated conductances in motoneurons were observed by 12 hours of blockade, but not changes in synaptic strength. It was unclear whether changes in voltage-gated conductances were observed by 2 hours of blockade when the recovery actually begins. Further, compensatory changes in voltage-gated conductances were not observed following glutamatergic blockade where embryonic movements were blocked but then recovered in a similar manner to GABAergic blockade. In this study, we discover a mechanism for homeostatic recovery in these first hours of neurotransmitter receptor blockade. In the first 6 hours of GABAergic or glutamatergic blockade there was a clear depolarization of resting membrane potential in both motoneurons and interneurons. These changes reduced action potential threshold and were mainly observed in the continued presence of the antagonist. Therefore, it appears that fast changes in resting membrane potential represent a key fast homeostatic mechanism for the maintenance of network activity in the living embryonic nervous system.

**Significance:** Homeostatic plasticity represents a set of mechanisms that act to recover cellular or network activity following a challenge to that activity and is thought to be critical for the developmental construction of the nervous system. The chick embryo afforded us the opportunity to observe in a living developing system the timing of the homeostatic recovery of network activity following 2 distinct perturbations. Because of this advantage, we have identified a novel homeostatic mechanism that actually occurs as the network recovers and is therefore likely to contribute to nervous system homeostasis. We found that a depolarization of the resting membrane potential in the first hour of the perturbations enhances excitability and supports the recovery of embryonic spinal network activity.

## Introduction

Recent work has focused on the mechanisms that allow networks to homeostatically maintain their activity levels in the face of various perturbations (1–3). Typically, activity is altered for 24 hours or more and compensatory changes in intrinsic cellular excitability and/or synaptic strength (synaptic scaling) are observed following the perturbation. While most of the work has been carried out *in vitro*, homeostatic mechanisms have also been observed *in vivo* in the spinal cord (4–6), hippocampal (7), auditory (8) and visual systems (9, 10). We have studied homeostatic plasticity in the chick embryo spinal cord, which expresses a spontaneously occurring network activity (SNA) that drives embryonic movements (11, 12). SNA likely occurs in all developing circuits shortly after synaptic connections form. In the embryonic spinal cord this activity is a consequence of the highly excitable nature of the nascent synaptic circuit where GABAergic neurotransmission is depolarizing and excitatory during early development (11–16). Spinal SNA is known to be important in motoneuron axonal pathfinding (17), and for proper muscle and joint development (18–23).

The embryonic spinal cord provides an exceptional model of homeostasis. The isolated embryonic spinal preparation exhibits SNA *in vitro*, which is transiently blocked by either glutamatergic or GABAergic receptor (GABAR) antagonists, but within hours is homeostatically restored in the presence of that antagonist (24, 25). As in the isolated cord *in vitro*, SNA-generated embryonic movements *in vivo* also exhibit a homeostatic recovery following blockade of either of the main excitatory neurotransmitter receptors. When GABA_A_ or glutamate receptor antagonists were injected into the egg at embryonic day 8 (E8), SNA-driven embryonic movements were abolished for 1-2 hours, but then homeostatically recovered to control levels 12 hours after the onset of pharmacological blockade of either transmitter (26). This provided us with the opportunity to assess mechanisms that contribute to the homeostasis of activity in the living system as it actually recovers. Although we initially thought that similar mechanisms would drive the recovery of embryonic activity following injection of either antagonist, we determined that compensatory changes in intrinsic excitability and synaptic scaling were only observed following blockade of GABA_A_ receptors, not glutamate receptors. Further, scaling was not observed by the time SNA-generated movements had fully recovered (12 hour GABAergic block). While compensatory changes in voltage-gated conductances were observed by this 12 hour time point, it was unknown if they occurred by 2 hours when the movements first start to recover (5), or if other mechanisms may be in place to initiate this recovery. In addition, the recovery of embryonic movements following glutamatergic blockade was very similar to that after GABAR blockade, but the mechanisms that drove this recovery remained completely unclear.

We had not examined the possibility that compensatory changes in cell excitability and/or scaling were occurring at the onset and throughout the recovery process in motoneurons. In fact, very few studies have compared the expression of presumptive homeostatic mechanisms with the timing of the homeostatic recovery of activity, yet we would expect that some of these mechanisms would be expressed at the very onset of the recovery process. Further, we had only focused on motoneurons in these previous studies, and had little knowledge about compensations that may be occurring in the interneurons that contribute to the drive of SNA. Therefore, we set out to identify the mechanisms that are expressed during the actual period of homeostatic recovery of embryonic activity *in vivo*. We find a previously unrecognized mechanism of homeostatic intrinsic plasticity where fast changes in resting membrane potential (RMP) bring both interneurons and motoneurons closer to action potential threshold. The results suggest that compensatory changes in RMP facilitate the homeostatic recovery of activity during glutamatergic or GABAergic blockade in the living embryo.

## Methods

### Dissection

*E*mbryonic day 10 (E10 or stage 36 (27)) chick spinal cords were dissected under cooled (15°C) Tyrode’s solution containing the following (in mM): 139 NaCl, 12 D-glucose, 17 NaHCO_3_, 3 KCl, 1 MgCl_2_, and 3 CaCl_2_; constantly bubbled with a mixture of 95% O2-5% CO_2_ to maintain oxygenation and pH around 7.3 (for a full description, see (4)). After the dissection, the cord was allowed to recover for at least 6 hrs in Tyrode’s solution at 18°C. The cord was then transferred to a recording chamber and continuously perfused with Tyrode’s solution heated to 27°C to allow for the expression of bouts of SNA with a consistent frequency.

### Electrophysiology

Whole-cell current clamp recordings were made from spinal motoneurons localized in lumbosacral segments 1-3 and were identified by their lateral position in the ventral cord. Recordings were also made from interneurons in the same segments but these were identified by their more medial position in the ventral cord. Patch Clamp tight seals (>2 GΩ) were obtained using electrodes pulled from thin-walled borosilicate glass (World Precision Instruments, Inc) in two stages, using a P-87 Flaming/Brown micropipette puller (Sutter Instruments) to obtain resistances between 5 and 10 MΩ. Once whole-cell configuration was achieved, voltage clamp was maintained for a period of 5 minutes to allow stabilization before switching to current clamp configuration. In some cases, whole cell voltage clamp recordings were also made from motoneurons and interneurons in order to acquire miniature postsynaptic currents (mPSCs) and these recordings were started after the 5 minute stabilization period. Series resistance during recording varied from 15 to 20 MΩ among different neurons and was not compensated. AMPA and GABA mPSCs were separated by their decay kinetics as described previously (Gonzalez-Islas and Wenner, 2006). The mPSCs were acquired on an Axopatch 200B patch clamp amplifier (Molecular Devices), digitized on-line using PClamp 10 (Molecular Devices), and analyzed using Minianalysis software (Synaptosoft). Bar charts and associated average values were obtained by determining an average mPSC amplitude for each cell (variable number of mPSCs/cell, 5pA cutoff), and then calculating the average of all cells. Recordings in current clamp were terminated whenever significant increases in input resistance (>20%) occurred. For voltage clamp experiments, recordings were terminated whenever significant increases in series resistance (> 20%) occurred. Both current and voltage clamp recordings were acquired using an AxoPatch 200B amplifier controlled by pClamp 10.1 software (Molecular Devices). The intracellular patch solution for both current and voltage clamp recordings contained the following (in mM): 5 NaCl, 100 K-gluconate, 36 KCl, 10 HEPES, 1.1 EGTA, 1 MgCl_2_, 0.1 CaCl_2_, 1 Na_2_ATP, and 0.1 MgGTP; pipette solution osmolarity was between 280 and 300 mOsm, and pH was adjusted to 7.3 with KOH. Standard extracellular recording solution was Tyrode’s solution (see above), constantly bubbled with a mixture of 95% O2-5% CO_2_. For embryos treated with saline or gabazine *in ovo*, a step protocol (1s, in 1pA increments) was employed to obtain rheobase, threshold voltage, and f-I relationships. To measure short-term changes in rheobase and threshold voltage following *in vitro* application of gabazine, a ramp protocol (from 0 to 200 pA; 1.2 s) was used (Figure 1A).

**Figure 1.**
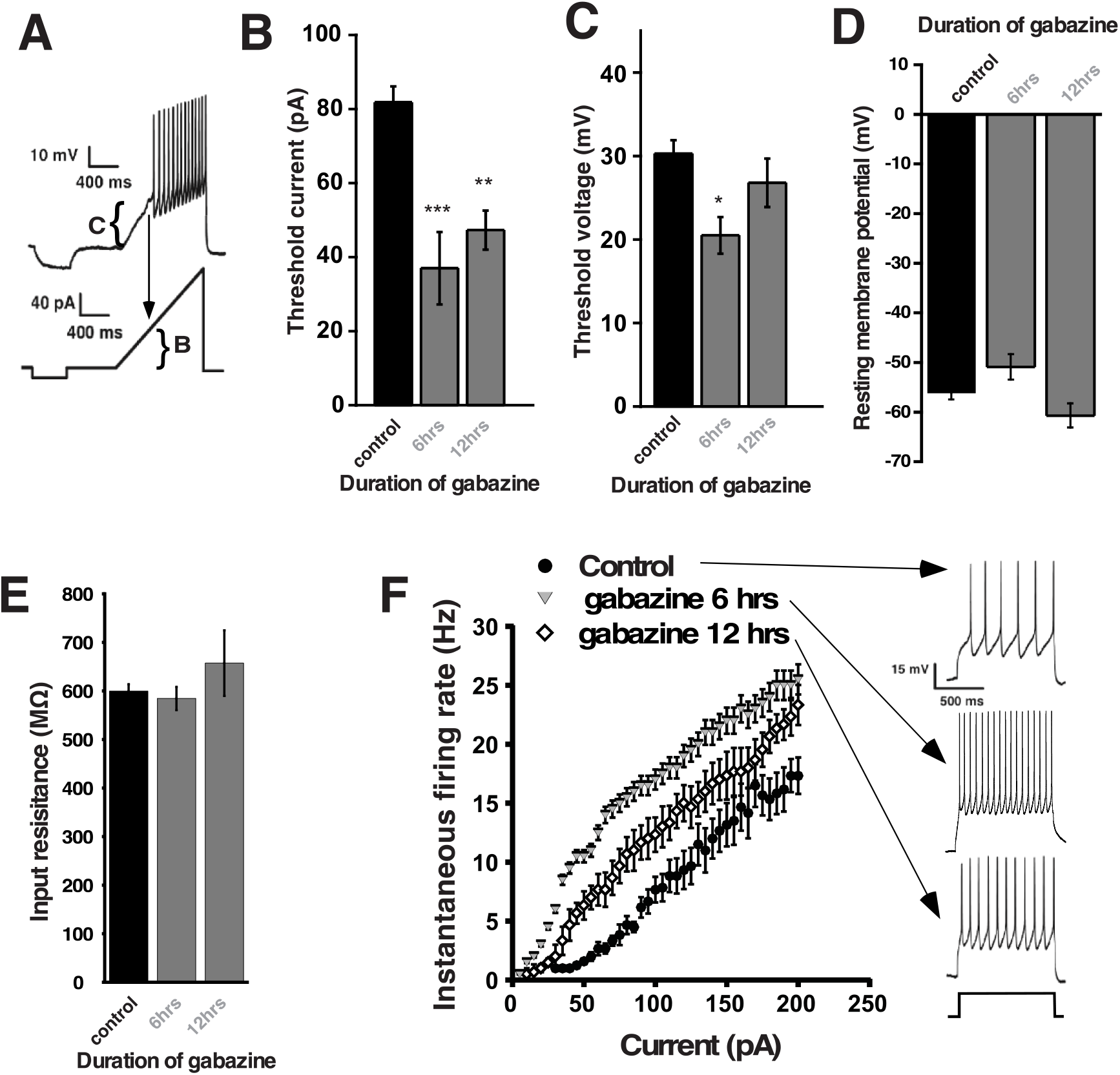
Changes in motoneuron excitability observed after chronic *in vivo* gabazine treatments. **A**) Motoneuron excitability was measured in isolated embryonic spinal cords by whole cell current clamp using a ramp protocol from 0 to 200 pA in 1.2 sec to assess rheobase current and voltage threshold. Representative spikes (top trace) evoked by somatic current injection (ramp protocol; bottom trace). Rheobase current (**B**) or threshold voltage (V_T_ –V_m_) (**C)** was obtained as the minimum current necessary to evoke a spike in the recorded motoneuron. Bar plotted summarizing the average rheobase current in control (n=12), and following chronic *in ovo* gabazine treatment (10 μM) for 6 hrs (n=11) and 12 hrs (n=9). No significant changes in resting membrane potential **(D**) or Input resistance (**E**) were found in the motoneurons recorded after chronic gabazine treatment at 6 or 12 hrs. **F**) Average f-I curves for control motoneurons (black circles; n=9), 6 hr gabazine treatment (grey triangles; n=6) or 12 hr gabazine treatment (open diamonds; n=9). Gabazine treatments shifted the average f-I curve to the left. The arrows point to representatives traces for each condition evoked by current steps of 100 pA. * P<0.05; ** P<0.01 and ***P<0.005.

### In ovo drug injections

A window in the shell of the egg was opened to allow monitoring of chick embryo movements and drug application 6 or 12 hrs before isolating the spinal cord at E10. 50 μl of a 10 mM gabazine solution was applied onto the chorioallantoic membrane of the chick embryo to a final concentration of 10µM, assuming a 50 ml egg volume.

### Immunoblots

The ventral half of the lumbosacral spinal cords were homogenized in RIPA buffer containing protease and phosphatase inhibitors. Samples were then centrifuged to remove cell debris. Protein concentration was quantitated using BCA reagent (Pierce). Samples were separated on 4–15% SDS-PAGE, and blotted to a nitrocellulose membrane. The primary antibodies against Nav 1.2 and Kv 4.2 and were from Alomone Labs. The blots were visualized by ECL chemiluminescence (GE Healthcare). Blots were done in duplicate, and each sample represents a lysate of 4-5 different cords, and were normalized to actin.

### Drugs

TTX and Gabazine were purchased from Tocris Cookson; All other chemicals and drugs were purchased from Sigma-Aldrich.

### Statistics

Data are expressed as mean ± SE. Statistical analysis of cellular excitability parameters was performed using ANOVA followed by Tukey’s post hoc test for multiple comparisons for normally distributed data and Kruskal-Wallis method followed by a post-hoc Dunn test for data that was not normally distributed. For statistical assessment of mPSC amplitude we used a student t-test for normally distributed data, and Mann-Whitney test for data that was not normally distributed. Throughout the manuscript * refers to p ≤ 0.05, ** refers to p ≤ 0.01, and *** refers to p ≤ 0.001.

## Results

### Changes in motoneuron excitability observed during the homeostatic recovery of SNA

In the embryonic spinal cord episodes of SNA are triggered through spiking of motoneurons that activate Renshaw cells (class of interneuron, (28)), which then recruit the rest of the network (29). Given the importance and accessibility of motoneurons in the initiation of SNA we have previously examined compensatory changes following GABAergic blockade in this cell type. In those studies we have shown that spinal motoneuron voltage-gated Na^+^ and K^+^ channel currents were altered following 12 hours of GABAergic blockade, after embryonic movements have homeostatically recovered (5). In order to determine if these changes actually contribute to the recovery, we assessed cellular excitability in motoneurons during the period that the SNA-driven movements were actually recovering, but before complete recovery was achieved. First, we tested whether cellular excitability had increased during the period that movements were in the process of homeostatically recovering, following 6 hours of gabazine treatment (10µM) *in ovo*. We isolated the spinal cord following treatment, and recorded whole cell in current clamp from motoneurons no longer in the presence of gabazine. We assessed threshold current (rheobase) and the voltage deflection produced by this current (threshold voltage, Figure 1, Table 1). Threshold current and voltage was reduced, compared to controls. As observed previously, threshold current was reduced following 12 hours of gabazine treatment *in ovo* (Figure 1). We also assessed excitability by giving current steps and plotting this against firing frequency after either 6 or 12 hours of gabazine treatment (Figure 1F). We saw a very strong shift toward higher excitability in the FI curve at the 6 hour time point, which then moved back toward pre-drug values following 12 hour treatment, although cells still showed a heightened excitability compared to controls. We did not observe changes in resting membrane potential or input resistance (Figure 1D, E).

One advantage of the earlier experiments was that the perturbation was carried out *in vivo*. Unfortunately, in order to measure cellular excitability the cord must be isolated and was given several hours to recover in the absence of gabazine before we could make excitability measurements. Such a process could itself alter the excitability of the cells. Therefore, in addition to the *in vivo* perturbations, we wanted to assess cellular excitability changes in the isolated cord *in vitro* as SNA was recovering, but still in the presence of gabazine. Control spinal cords were isolated and expressed episodes of SNA *in vitro* at a fairly constant frequency and then gabazine (10µM) was added to bath and whole cell recording of motoneurons were obtained in the presence of gabazine at different points after the addition of the GABA_A_ antagonist. Following *in vitro* 6 hour (4-6 hours) gabazine treatment, motoneuron cellular excitability was increased (Figure 2A-B) similarly to that of the 6 hour *in ovo* treatment (Figure 1B-C). We also measured excitability in spinal motoneurons following shorter periods of *in vitro* gabazine exposure. Like the 6 hour treatment, we found that motoneuron cellular excitability was also increased in the first 2 hours and from 2-4 hours of *in vitro* exposure to gabazine (Figure 2A-B). We saw that changes in threshold voltage were between 10-17mV, and that a depolarization the RMP could account for ∼5mV of this change in threshold voltage. RMP was reduced at 0-2, 2-4, and 4-6 hour time points, but did not reach significance in any of these periods. On the other hand if we compared control RMP to gabazine-treated at all time points combined, then there was a significant depolarization of RMP (p=0.03, Figure 2C). We saw no change in input resistance in any of the conditions (Figure 2D). These results suggest that compensatory changes in motoneuron excitability occurs very quickly and therefore could contribute to the recovery of SNA, and part of this shift in threshold was due to a change in the RMP.

**Figure 2.**
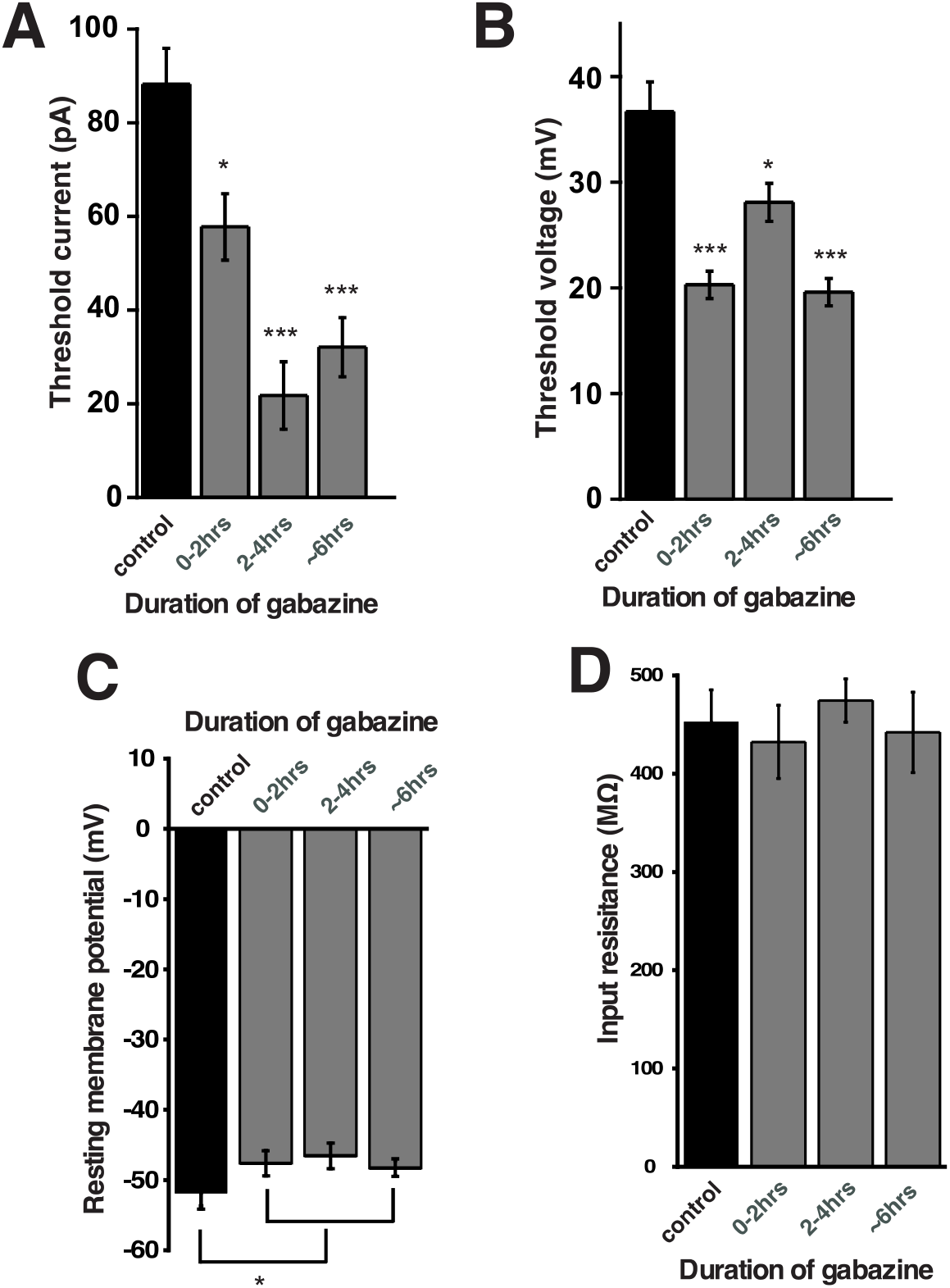
Changes in motoneuron excitability observed *in vitro* before and after the blockade of GABA_A_ receptors. Motoneuron excitability was measured in isolated spinal cords in the absence (control) or in the continuous presence of gabazine 10 μM at 3 different blockade periods: 0-2 hrs; 2-4 hrs and 4-6 hrs **A**) Bar plot showing the average value of the rheobase current in control (n=13; black bar); 0-2 (n=13); 2-4 (n=9) and 4-6 (n=10) hrs in gabazine. **B**) Summary of the values of the average spike threshold (V_T_ - V_m_) during the same intervals. No significant changes in the resting membrane potential (**C**) or input resistance (**D**) were found in separate periods, but RMP was different when 3 gabazine periods combined and compared to control. A ramp protocol from 0 to 200 pA in 1.2 s was used to assess rheobase current and voltage threshold. * P<0.05; ** P<0.01 and ***P<0.005.

### Changes in interneuron excitability occur during the recovery of SNA following GABAergic blockade

We wanted to determine whether these increases in cellular excitability were only occurring in motoneurons, or whether this was a more general phenomenon that extends to the rest of the developing motor circuitry. Thus, we assessed the possibility that spinal interneurons also increased cell excitability following gabazine treatment and could therefore contribute to the homeostatic recovery of SNA. Spinal neurons were targeted in the more medial positions of the cord. We found that, like motoneurons, interneurons had reduced threshold current and voltage (increased cellular excitability) following both 6 and 12 hours of gabazine treatment *in ovo* (Figure 3A-B, Table 1). Similar increases in cellular excitability were also observed in interneurons from isolated cords that were treated with bath application of gabazine for 0-2, 2-4, and 4-6hrs *in vitro* (Figure 3A-B, Table 1). Because we were recording from diverse classes of spinal neurons, the results suggest that various cell types alter their cellular intrinsic excitability and contribute to the recovery of activity following GABAergic blockade. Importantly, interneuron RMP was significantly depolarized at each of the time points (Figure 3D). The threshold voltage after gabazine treatment (*in vitro* or *in vivo*) at any of the times was reduced by ∼10-18mV. A depolarized RMP could account for the change in threshold voltage following the *in vitro* gabazine studies (11-17mV), but only partially could explain the results following 6 hour *in vivo* gabazine treatment (Figure 3B, D). No changes were observed in input resistance in any condition. Therefore, interneurons increased their cellular excitability after GABAergic blockade similarly to motoneurons. However, interneurons achieved the increase in excitability following *in vitro* gabazine solely through a depolarized RMP, unlike motoneurons.

**Figure 3.**
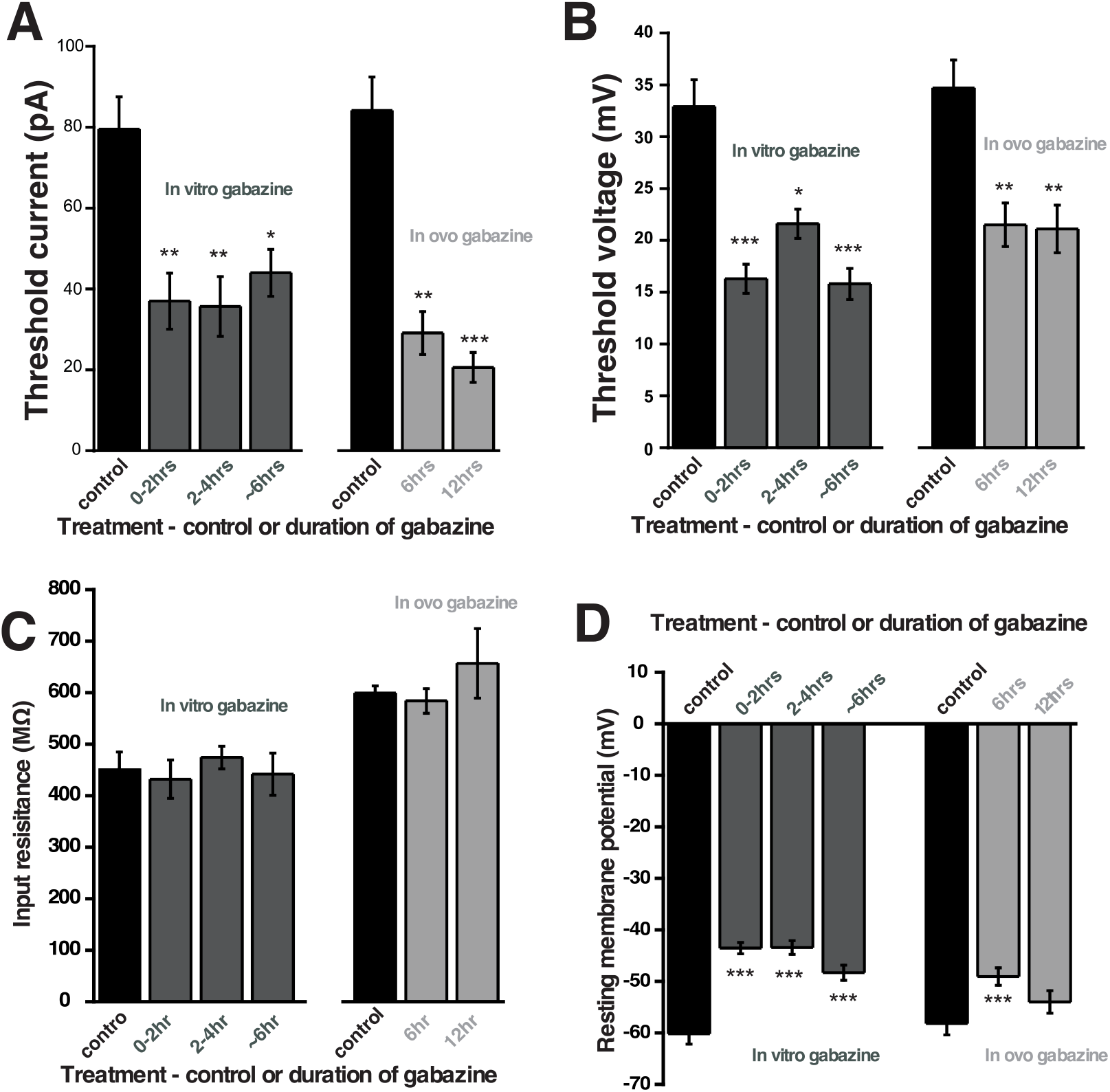
Interneuron excitability increased following *in vivo* and *in vitro* GABAergic blockade. Interneuron excitability was measured in isolated spinal cords after *in ovo* gabazine treatment (right) or acutely *in vitro* for different periods (left) during the period of recovery of SNA: *In vitro*, in the presence of gabazine 10 μM, at intervals of 0-2 hrs (n=14), 2-4 hrs (n=9) and 4-6 hrs (n=6) and after chronic treatment with gabazine 10 μM *in ovo* for 6 (n=12) or 12 hrs (n=9) in the absence of gabazine. Bar charts show average values for spike threshold current (Rheobase) (**A**), spike threshold voltage (V_T_ - V_m_) **(B**), input resistance **(C**), or resting membrane potential **(D**). A ramp protocol from 0 to 200 pA in 1.2 s was used to assess rheobase current and voltage threshold. * P<0.05; ** P<0.01 and ***P<0.005.

Since the changes in cellular excitability following GABAR blockade appear to be expressed across multiple cell types throughout much of the cord, we ran Western blots of isolated spinal cords (ventral half) and assessed 2 of the voltage-gated channels that we expected could mediate this process. We reported in an earlier study (5) that gabazine-induced changes were observed in voltage-gated Na^+^ and K^+^ channels. In that study we saw TTX-sensitive voltage-gated Na^+^ channel currents were increased. Therefore, we assessed the levels of Nav1.2, an alpha subunit of the voltage-gated Na^+^ channel, which had been shown to be expressed early in the development of the embryonic chick (30). We found that following 12 hour gabazine treatment *in ovo*, Nav1.2 expression was increased (172.4±14.8%, p ≤ 0.05), but not after 6 hour treatment (105.3 ± 5.3%, Figure 4). Further, we had observed that currents of the A-type transiently-activated K^+^ channel (I_A_) and the calcium-dependent K^+^ channel (I_K(Ca)_) were both decreased following gabazine treatment (5). Here, we show that expression of Kv4.2 (mediates the A-type K^+^ channel in chick embryo (31)) is down regulated following 12 hour gabazine treatment (54.5 ± 2.5%, p ≤ 0.05), but not after 6 hours (104.2 ± 17.2%, Figure 4). Together the results show that cellular excitability is altered during and after the homeostatic recovery of SNA, and that changes in total voltage-gated channel expression do not contribute to recovery of excitability at short periods but could contribute at later stages of the recovery.

**Figure 4.**
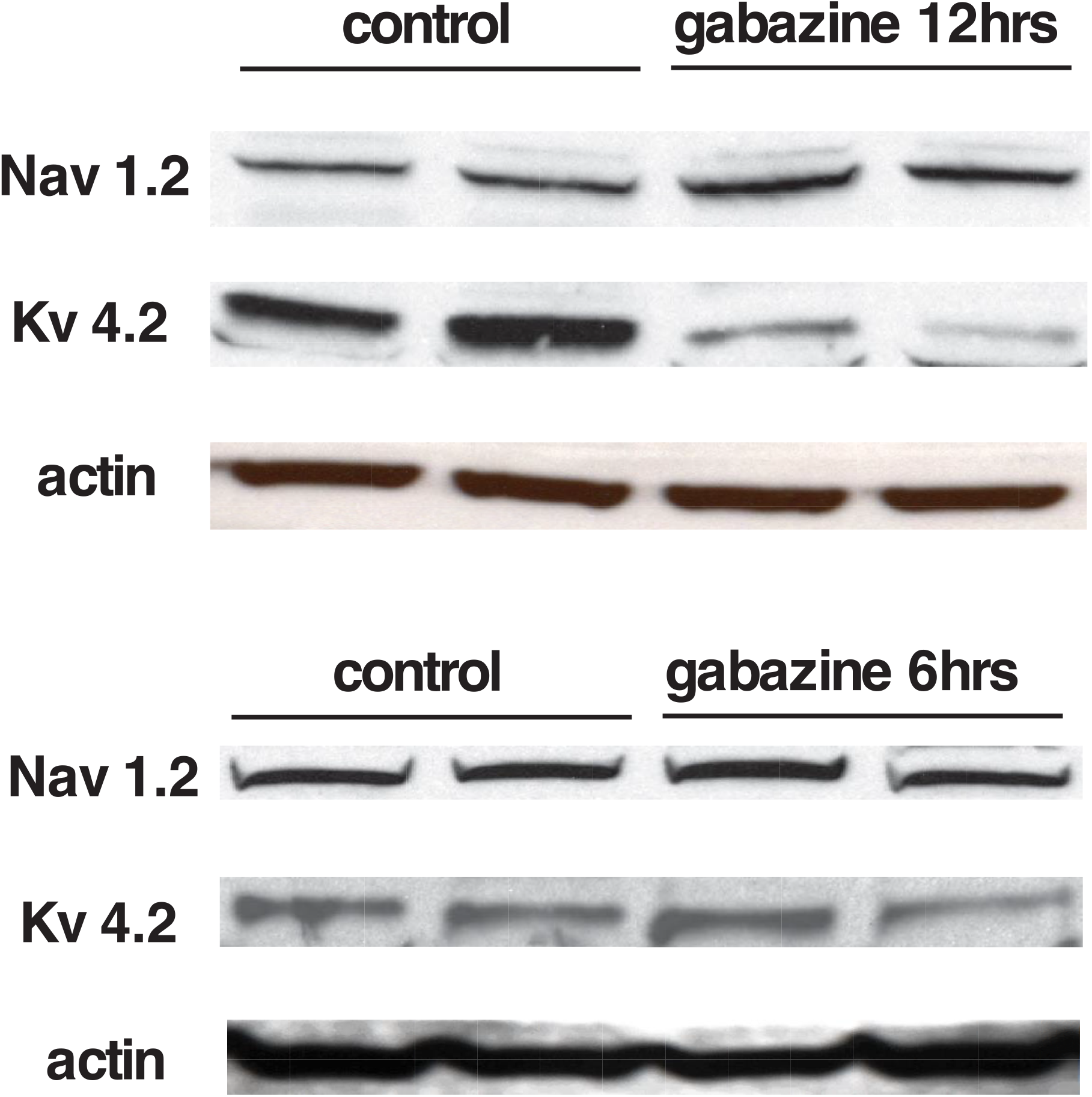
Changes in voltage-gated channel expression following *in ovo* gabazine treatment. Western blots showing changes in the expression of voltage-dependent Na^+^ channels (Nav1.2) and inactivating K^+^ channels (Kv4.2) in the chick embryo spinal cord following 12 hrs **(A)** or 6 hrs **(B)** of GABA_A_ receptor blockade *in ovo*.

### The trigger for changes in RMP was distinct from the homeostatic mechanisms expressed after GABAergic blockade

Following GABA_A_R blockade compensatory changes in synaptic strength (scaling) were not observed until 48 hours, but voltage-gated conductance changes were triggered by 12 hours (5, 26). More recently we have determined that simply reducing GABA_A_R activation due to spontaneous miniature release of GABA vesicles (spontaneous GABAergic transmission) was sufficient to trigger upscaling (32). Therefore, we tested the possibility that the fast changes in cellular excitability observed in the current study were also mediated by reduced spontaneous miniature GABAergic neurotransmission. As in previous studies we took advantage of the nicotinic antagonist DHβE, which reduces spontaneous GABA release, to test this possibility. Whole cell recordings from motoneurons were obtained before and 2 hours after DHβE application *in vitro*. We did not find any differences in cellular excitability following reduction of GABA release by DHβE (Figure 5). Therefore, unlike the trigger for synaptic scaling, the compensatory changes in cellular excitability in the first hours of GABAergic blockade were not mediated by changes in spontaneous GABAergic transmission.

**Figure 5.**
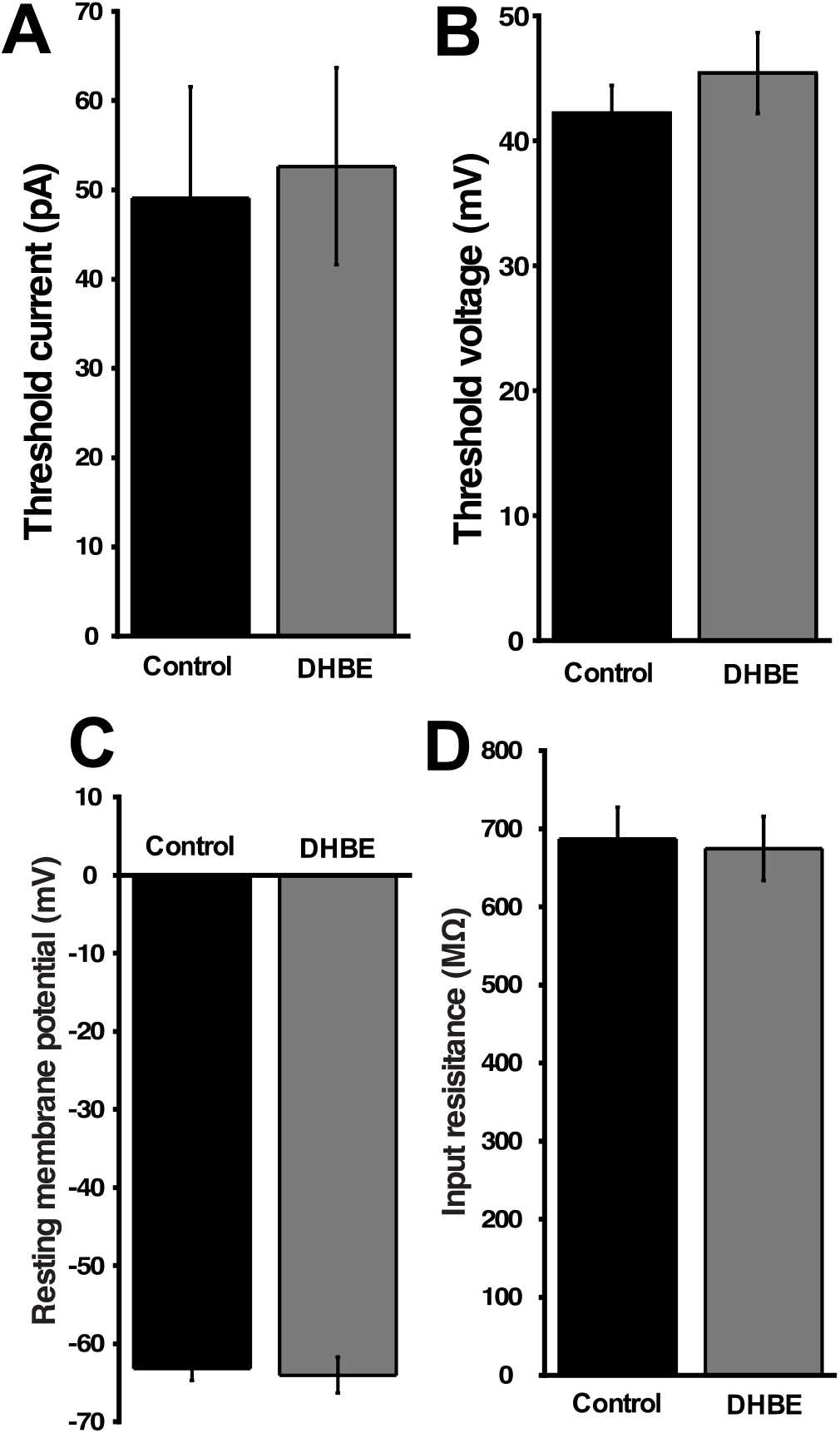
Chronic reductions in quantal GABA release that mediate synaptic upscaling do not trigger changes in motoneuron excitability. Whole cell current clamp recordings from motoneurons were obtained from isolated spinal cords before (control; black bars; n= 11) or after 2hrs of the nicotinic receptor antagonist (DHβE 10 μM, grey bars; n=9). Average rheobase current (**A**) average spike threshold voltage (V_T_ -V_m_) **(B)** RMP, and input resistance were no different different after two hours of DHβE compared to controls. A ramp protocol from 0 to 200 pA in 1.2 s was used to assess rheobase current and voltage threshold.

### Synaptic scaling does not contribute to the homeostatic recovery of SNA-generated movements

Previously we showed that AMPAergic and GABAergic upscaling were not observed in chick embryo motoneurons following 12 hour gabazine treatment *in ovo*. However, it remained possible that interneurons experienced scaling and contributed to the homeostatic recovery of SNA. We had never directly assessed mPSC scaling in interneurons previously, so it was possible that scaling in spinal interneurons occurred in the first hours of gabazine treatment. However, following 6hrs of gabazine treatment *in ovo* we found no increase in AMPA mPSC amplitude in interneurons (Figure 6A). We would not have expected GABAergic scaling to contribute to the recovery of movements as we were blocking GABA_A_ receptors. Regardless, we examined the possibility that GABAergic blockade triggered GABAergic scaling in interneurons as we previously found slight increases in interneuronal Cl^-^, which mediated GABAergic scaling following 24 hours of gabazine treatment *in ovo* (33, 34). We did not see any change in GABA mPSC amplitude in interneurons following 6hr gabazine treatment *in ovo* (Figure 6A). The results show that 6 hour gabazine treatment did not trigger either AMPAergic or GABAergic scaling in interneurons, and so would not contribute to the homeostatic recovery of activity levels.

**Figure 6.**
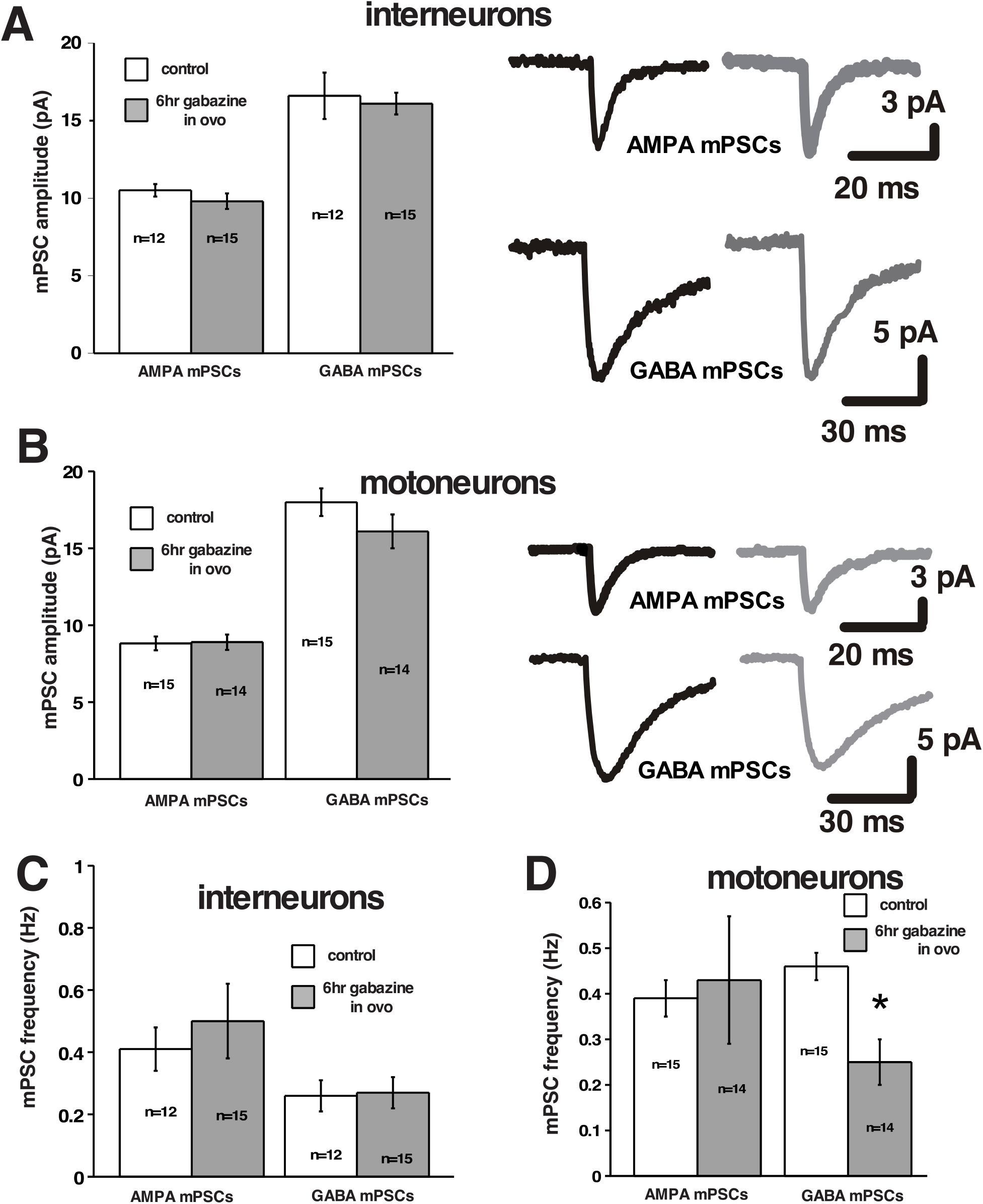
Synaptic scaling does not contribute to the homeostatic recovery of SNA-generated movements. **(A)** Bar plot showing AMPAergic and GABAergic miniature postsynaptic current amplitudes (mPSCs) from embryonic spinal interneurons in control (white bar; n=12) and after 6 hrs of chronic treatment with 10 μM gabazine (grey bar; n=15). **(B)** Bar plot showing AMPAergic and GABAergic miniature postsynaptic current amplitudes (mPSCs) from embryonic spinal motoneurons in control (white bar; n=12) and after 6 hrs of chronic treatment with gabazine 10 μM (grey bar; n=15). **(C, D)** Bar chart showing AMPAergic and GABAergic miniature postsynaptic current frequencies from embryonic spinal interneurons **(C)** or motoneurons **(D)** in control (white bar) or after 6 hrs of chronic treatment with gabazine 10 μM (grey bar).

We have reported that scaling is not triggered in motoneurons after 12 hours of gabazine treatment, but had not checked if it was transiently expressed in the first hours as the network recovered as has been described in cultured neurons (35, 36). We therefore tested for AMPAergic and GABAergic scaling in motoneurons following 6 hours of gabazine treatment *in ovo*. We found that AMPAergic and GABAergic mPSC amplitude was no different than controls (Figure 6B). These findings showed that synaptic scaling of AMPAergic mPSCs in different spinal populations could not have contributed to the recovery of SNA or the movements it drives following GABAergic blockade. Finally, there was no compensatory increase in mPSC frequency in interneurons (Figure 6C), or motoneurons (Figure 6D). In fact, we found a significant reduction in GABAergic mPSC frequency.

### Recovery of embryonic movements following glutamatergic blockade is mediated by fast changes in resting membrane potential

Previously, we had carried out experiments where a similar homeostatic recovery of embryonic movements was observed following glutamatergic blockade, instead of GABAergic blockade. Following GABAergic or glutamatergic blockade movements recovered in around 12 hours, but after 12 hours of glutamatergic blockade we did not see synaptic scaling or homeostatic changes in intrinsic excitability. However, we never looked at earlier time points in the presence of glutamatergic antagonists to see if there were changes in intrinsic excitability. Therefore, we isolated spinal cords and tested for compensatory changes in intrinsic excitability from 0-6 hours of CNQX (20µM)/APV (50µM) application *in vitro* in the continued presence of the antagonists. We observed that motoneurons did indeed express reductions in threshold current and voltage (15-20mV) in the first 6 hours of drug application (Figure 7A-B). As following GABAergic blockade, the reduction in threshold voltage was reasonably well matched by a similar depolarization of resting membrane potential (12-14mV), and no change in input resistance was seen (Figure 7C-D). Similarly interneurons showed the same response to glutamatergic blockade, a fast increase in threshold voltage (∼15mV) mediated by a depolarization of the RMP (17mV) (Figure 8). The results suggest fast homeostatic changes in membrane potential can largely account for changes in threshold voltage for motoneurons and interneurons following glutamatergic blockade. Together, the results suggest that homeostatic changes in RMP likely contribute to the recovery of embryonic movements during either GABAergic or glutamatergic blockade.

**Figure 7.**
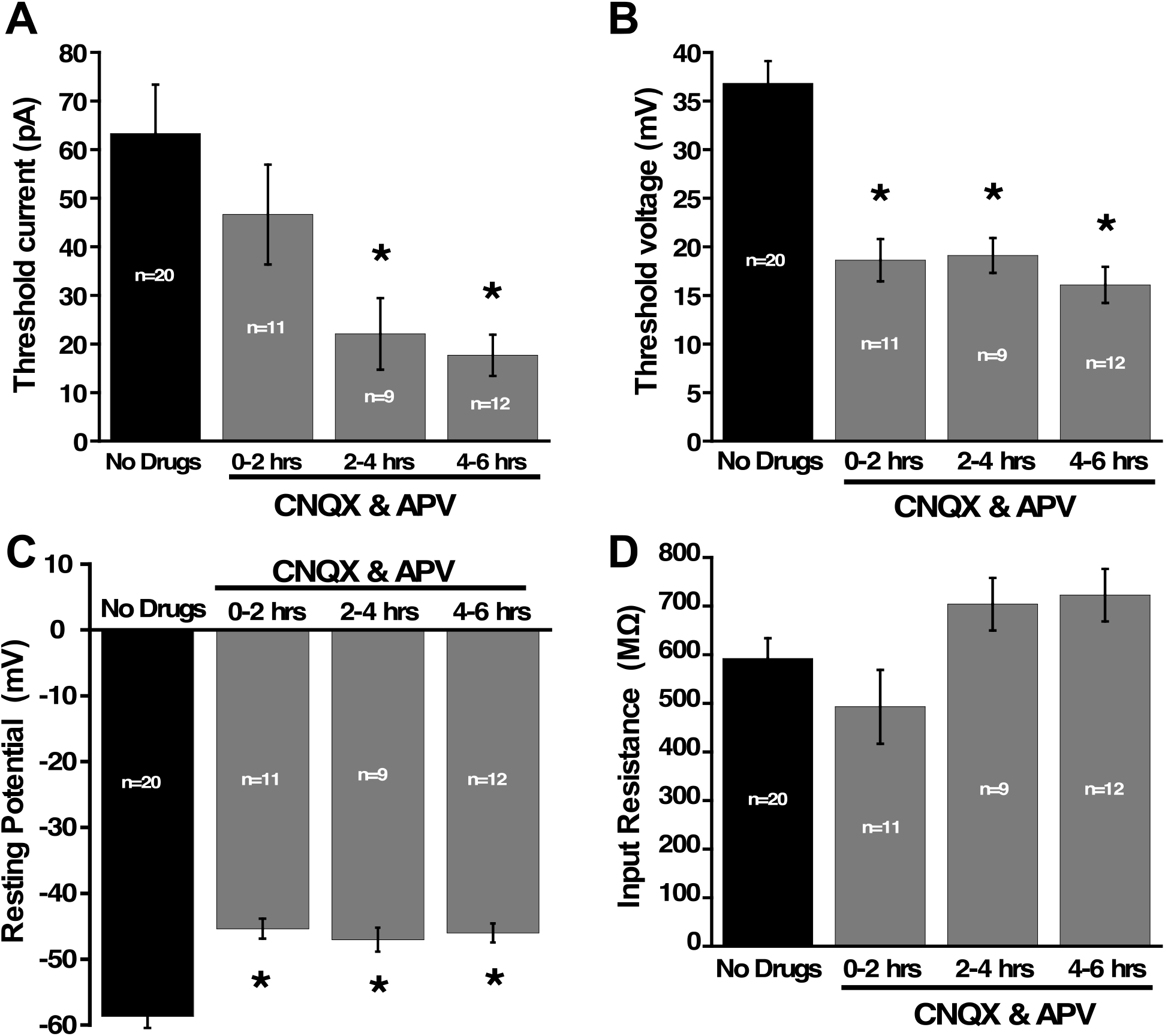
Changes in motoneuron excitability observed during blockade of glutamatergic receptors is mediated by fast changes in resting membrane potential. Whole cell current clamp recordings from motoneurons were obtained from isolated spinal cords before and during glutamatergic blockade (CNQX 10 μM and APV 25 μM). Rheobase current **(A)** and spike threshold voltage (V_T_ -V_m_) **(B)** were measured through a ramp protocol from 0 to 200 pA in 1.2 seconds in the absence or under the presence of glutamatergic blockade at 3 different durations of blockade: 0-2 hrs (n=11); 2-4 hrs (n=9) and 4-6 hrs (n=12). **C**) Increased excitability in motoneurons was due to a depolarization of the resting membrane potential occurring in the motoneurons after the addition of glutamatergic receptor blockers. **D**) No significant changes in input resistance were observed in the recorded motoneurons over the same period of time. * P<0.05.

**Figure 8.**
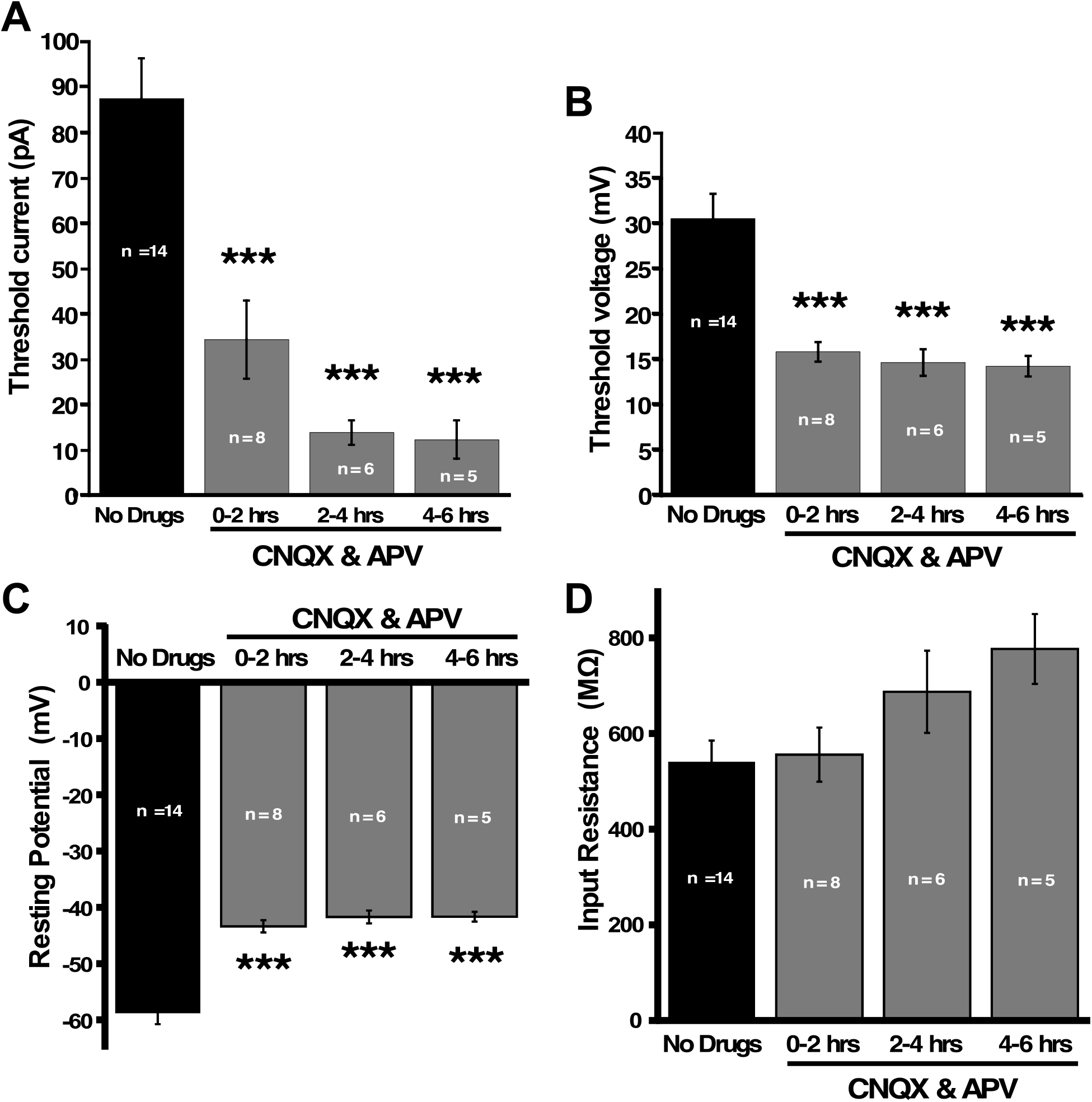
Changes in interneuron excitability observed during blockade of glutamatergic receptors is mediated by fast changes in resting membrane potential. Whole cell current clamp recordings from motoneurons were obtained from isolated spinal cords before and during glutamatergic blockade (CNQX 10 μM and APV 25 μM). Rheobase current **(A)** and spike threshold voltage (V_T_ -V_m_) **(B)** were measured through a ramp protocol from 0 to 200 pA in 1.2 seconds in the absence or under the presence of glutamatergic blockade at 3 different durations of blockade: 0-2 hrs (n=8); 2-4 hrs (n=6) and 4-6 hrs (n=5). **C**) Increased excitability in interneurons was due to a depolarization of the resting membrane potential occurring in the interneurons after the addition of glutamatergic receptor blockers. **D**) No significant changes in input resistance were observed in the recorded interneurons over the same period of time. *** P<0.005.

## Discussion

### Homeostatic changes in RMP, synaptic strength, and voltage-gated conductances and their contribution to the recovery of embryonic activity

Synaptic scaling was not expressed at 6 hours in motoneurons or interneurons (Figure 6), and therefore this form of homeostatic plasticity does not appear to be involved in the recovery of embryonic spinal activity, consistent with our previous study (26). While scaling does not appear to mediate the recovery in the embryonic spinal cord, it appears to influence recovery of spiking activity in the cortex following *in vivo* sensory deprivation (37, 38).

The initial homeostatic changes in RMP (<2hrs) occur well before synaptic scaling (24-48 hrs) in the chick embryo (26, 33), and are observed in motoneurons and interneurons during either GABAergic or glutamatergic blockade. Therefore, this form of fast homeostatic plasticity is very likely to contribute to the recovery of embryonic movements and the SNA that drives this activity. On the other hand, there were also compensatory changes in threshold voltage in motoneurons and interneurons during GABAergic blockade *in vitro* and following 6 and 12 hours of transmitter blockade *in vivo* that were not completely explained by changes in RMP. These changes were likely to be similar to the previously described changes in voltage-gated conductances associated with homeostatic intrinsic plasticity following 12 hours of GABAergic blockade (5). Changes in protein levels of 2 previously implicated voltage-gated ion channels were only observed at 12 hours, although not at 6 hours of *in vivo* gabazine treatment (Figure 4). Therefore, these gabazine-induced changes in intrinsic plasticity could contribute to the recovery of activity by 12 hours along with changes in RMP. Interestingly, following glutamatergic blockade, the changes in threshold voltage (∼15mV) could be completely explained by changes in RMP (∼15mV, Figure 7-8). The recovery of embryonic movements following GABAergic and glutamatergic blockade are temporally very similar, however long-term changes in voltage-gated conductances *in ovo* (5) or in the first 6 hours *in vitro* (Figure 7-8) were only observed following GABAergic blockade. This is consistent with the idea that changes in voltage-gated conductances are not necessary for the recovery and would place compensatory changes in RMP as a more important mechanism of homeostatic recovery of SNA in this developing circuitry.

### Homeostatic perturbations and RMP

No changes in RMP were observed in two of the earliest studies on homeostatic plasticity where different strategies were used to chronically block spiking for days (TTX, CNQX, or cell isolation) - in rat cortical cultures (39) or the stomatogastric neurons of the lobster (40). Since these early studies, several other homeostatic experiments have been performed where spiking or neurotransmission were chronically blocked to trigger homeostatic synaptic or intrinsic plasticity and no changes in RMP were observed. These studies have been carried out using various perturbations (TTX, TTX/APV, CNQX, NBQX, gabazine, bicuculline, cell isolation) in rat cortical cultures (39), mouse cortical cultures (41), and in the embryonic spinal cord of the chick embryo *in* vivo (4, 5, 26). On the other hand, none of these studies followed the RMP before and immediately after the perturbation as we have done in the current study. Therefore, changes in RMP may have occurred in these previous studies but over the duration and/or after removal of the perturbation no change was detected. An exception to this was described following a fairly severe perturbation (2 week exposure to 15mM KCl), where a homeostatic hyperpolarization of RMP was observed in cultured hippocampal neurons (42).

### Conductances that contribute to the RMP and the timing of homeostatic changes

Several different ion channels exhibit activation at subthreshold potentials and thus contribute to setting the RMP including multiple kinds of K^+^ channels (Ia, Ikir, Ileak) (43, 44), hyperpolarization-activated cationic channels (Ih) (45, 46), low-threshold calcium channels (47), persistent sodium currents (NaPIC) (48, 49), and leak sodium channels (NALCN) (50, 51). In addition, ongoing synaptic conductances can also influence the RMP (52, 53). Previous work has shown that blocking GABARs by direct application of a GABA receptor antagonist onto a chick embryo spinal cord preparation causes an acute hyperpolarization in spinal neurons that can be as large as 10mV, suggesting a significant tonic GABAergic depolarizing current (54). The effect of acute application of GABAergic antagonist onto the cord (54) is in the opposite direction (hyperpolarizing) compared to the current studies finding that bath application of gabazine leads to a depolarization of RMP in the first hours of drug exposure. The current study is the only one we are aware of that follows RMP before and throughout the first hours of the perturbation, and may explain why this form of homeostatic intrinsic plasticity has not been previously reported.

### Mechanisms of homeostatic changes in RMP

What is a potential trigger for these homeostatic changes in RMP? Previously we have shown that homeostatic synaptic scaling is triggered following 48 hour block of GABAergic transmission (26) and compensatory changes in voltage-gated ion channel conductances by 12 hours of GABAR block (5). In fact, we have also shown that merely altering GABAR activation due to spontaneous release of GABA vesicles can fully trigger synaptic scaling. However, compensatory changes in RMP were not so reliant on GABAR activation. Fast changes in RMP are not triggered by altering the frequency of spontaneous vesicle-mediated GABAR activation (Figure 5). Further, these changes can also be triggered by reduced glutamatergic receptor activation where GABAR activation is intact. GABAergic or glutamatergic receptor blockade both trigger compensatory changes in RMP and exhibit a temporally similar recovery of movements. Therefore, the most straightforward explanation for the trigger of homeostatic changes in RMP would be the reduction in network activity caused by blocking either glutamatergic or GABAergic receptors. Compensatory changes in RMP driven by network activity would fit nicely with a direct feedback mechanism where activity is regulated by changes in activity.

There are two general mechanisms that could mediate the quick homeostatic changes in RMP. A commonly described mechanism underlying a change in RMP involves a change in some channel conductance (e.g. K^+^ channels). While this is certainly a possibility, we did not see a change in input resistance. Therefore, multiple channel conductances would need to change in an orchestrated manner such that total conductance remained unchanged.

A potentially more plausible mechanism for our observations would involve a change in the function of the Na^+^/K^+^ ATPase. Previous studies are consistent with this possibility. First, it has been shown that temperature influences spike frequency through adjustments in the resting membrane potential of invertebrate neurons, and this is mediated by an alteration of the electrogenic Na^+^/K^+^ ATPase (55–57). Next, work in the spinal cord of xenopus and neonatal mice, as well as in motoneurons of the fly larva, show that bursts of spiking activity expressed in these systems lead to an increase in intracellular Na^+^, that is necessary to activate an isoform of the Na^+^/K^+^ ATPase that is not active at baseline Na^+^ levels. The Na^+^-dependent activation of this Na^+^/K^+^ ATPase produces a hyperpolarizing current due to the electrogenic nature of the pump that has been called an ultra slow afterhyperpolarization (usAHP) (58–61). This hyperpolarizing current is maintained for up to a minute. SNA in the chick embryo spinal preparation experiences a very similar usAHP after episodes of SNA. Further, we have established that embryonic spinal neurons have very high Na^+^ concentrations at baseline (62). Therefore, it is possible that Na^+^ levels constitutively activate this Na^+^/K^+^ ATPase and when SNA is blocked for many minutes by glutamatergic or GABAergic antagonists, Na^+^ levels eventually are reduced to a point that pump activity is minimized and the hyperpolarizing current abates, thus depolarizing the RMP.

Alternatively, a downregulation of the Na^+^/K^+^ ATPase activity could produce the depolarization of RMP through a weakening of the K^+^ gradient. In fact, this has been described following chronic disuse of muscle fibers (63–65). Consistent with a plasticity of the Na^+^/K^+^ ATPase, it is clear that Na^+^, as well as Cl^-^, concentration is dynamically regulated during development in embryonic spinal motoneurons, and this is consistent with the idea that K^+^ will also be dynamically regulated during this period (33, 62, 66).

Most of our results appear to suggest that the changes in RMP are expressed transiently while the antagonists are in place, and once washed off the RMP returns to pre-drug levels. This kind of temporary perturbation might not permanently change the developmental trajectory of spinal neurons or their network. However, if this initial fast homeostatic mechanism does not recover activity levels or is maintained for long periods then other mechanisms may be triggered, which could alter the development of the spinal circuitry in a longer term manner. In some cases homeostatic changes in RMP may be sufficient and exist temporarily. This could obviate the need for other homeostatic mechanisms. Therefore, it will be important to see if compensatory changes in RMP trigger other forms of plasticity or are sufficient to recover activity by themselves, leaving the network relatively unperturbed.

## Acknowledgements

This research was supported by Grants from the National Institute of Neurological Disorders and Stroke to P. Wenner. We thank Drs. Mark Rich, Morten Raastad, Dobromila Pekala, and Pernille Bülow, Brendan O’Flaherty, Kun Lin for their valuable comments on the manuscript.

---

Cellular Excitability Measurements.

Cellular Excitability Measurements
| Condition | MN<br>Threshold<br>current (pA) | MN<br>Threshold<br>voltage (mV) | number<br>of cells | IN<br>Threshold<br>current (pA) | IN Threshold<br>voltage (mV) | Number of<br>cells |
| --- | --- | --- | --- | --- | --- | --- |
| Control in vitro | 88.2 ± 7.7 | 36.7 ± 2.8 | 13 | 79.5 ± 8.0 | 32.9 ± 2.6 | 12 |
| Gbz-treated<br>0-2hrs in vitro | 57.8 ± 7.1<br>p ≤ 0.05 | 20.3 ± 1.3<br>p ≤ 0.001 | 13 | 37.0 ± 6.9<br>p ≤ 0.01 | 16.3 ± 1.4<br>p ≤ 0.001 | 14 |
| Gbz-treated<br>2-4hrs in vitro | 21.8 ± 7.2<br>p ≤ 0.001 | 28.1 ± 1.8<br>p ≤ 0.05 | 9 | 35.7 ± 7.4<br>p ≤ 0.01 | 21.6 ± 1.4<br>p ≤ 0.05 | 9 |
| Gbz-treated<br>4-6hrs in vitro | 32.1 ± 6.3<br>p ≤ 0.001 | 19.6 ± 1.3<br>p ≤ 0.001 | 10 | 44.0 ± 5.8<br>p ≤ 0.05 | 15.8 ± 1.5<br>p ≤ 0.001 | 6 |
| Control in ovo | 81.8 ± 4.3 | 30.3 ± 1.6 | 12 | 84.1 ± 8.3 | 34.7 ± 2.7 | 10 |
| Gbz-treated<br>6hrs in vivo | 37.0 ± 9.8<br>p ≤ 0.001 | 20.5 ± 2.2<br>p ≤ 0.05 | 11 | 29.1 ± 5.3<br>p ≤ 0.01 | 21.5 ± 2.1<br>p ≤ 0.01 | 12 |
| Gbz-treated<br>12hrs in vivo | 47.3 ± 5.3<br>p ≤ 0.01 | 26.8 ± 2.9 | 9 | 20.6 ± 3.7<br>p ≤ 0.001 | 21.1 ± 2.3<br>p ≤ 0.01 | 9 |

